# EEG correlates of working memory performance in females

**DOI:** 10.1101/098301

**Authors:** Yuri G. Pavlov, Boris Kotchoubey

## Abstract

**Background:** The study investigates oscillatory brain activity during working memory (WM) tasks. The tasks employed varied in two dimensions. First, they differed in complexity from average to highly demanding. Second, we used two types of tasks, which required either only retention of stimulus set or retention and manipulation of the content. We expected to reveal EEG correlates of temporary storage and central executive components of WM and to assess their contribution to individual differences.

**Results:** Generally, as compared with the retention condition, manipulation of stimuli in WM was associated with distributed suppression of alpha1 activity and with the increase of the midline theta activity. Load and task dependent decrement of beta1 power was found during task performance. Beta2 power increased with the increasing WM load and did not significantly depend on the type of the task.

At the level of individual differences, we found that the high performance (HP) group was characterized by higher alpha rhythm power. The HP group demonstrated task-related increment of theta power in the left anterior area and a gradual increase of theta power at midline area. In contrast, the low performance (LP) group exhibited a drop of theta power in the most challenging condition. HP group was also characterized by stronger desynchronization of beta1 rhythm over the left posterior area in the manipulation condition. In this condition, beta2 power increased in the HP group over anterior areas, but in the LP group over posterior areas.

**Conclusions:** WM performance is accompanied by changes in EEG in a broad frequency range from theta to higher beta bands. The most pronounced differences in oscillatory activity between individuals with high and low WM performance can be observed in the most challenging WM task.

## Background

The ability to retain information in memory for a short period of time is critical for numerous cognitive tasks including planning, verbal competence, spatial orientation, mental manipulations of objects and many others [12–14].

According to Baddeley & Hitch’s [15] model, the structure of working memory (WM) consists of several components. One of them is responsible for temporary storage of information in modality-specific buffers. Another key component, the central executive, is considered to be a set of tools designed to maintain the active representation of memory trace, to control attention and to preserve the latter from interference caused by irrelevant stimuli [16,17].

A number of neuroimaging studies demonstrated that maintenance of information in WM engages a broad network of neural structures mostly including prefrontal cortex, parietal and temporal areas [13,18]. Whereas storage buffers represent information received from sensory inputs in posterior regions, the prefrontal cortex sustains and transforms this information and organizes executive processes of working memory [19]. Existing research highlights the importance of the fronto-parietal network activation in working memory processes, especially in high demanding tasks [20–24]. Apparently, individual differences in working memory capacity are also determined by fronto-parietal white matter connectivity [25].

Features of the processes presumed by Baddeley & Hitch’s model of WM cannot be characterized only by spatial distribution of brain activation. Qualitatively different information about these processes can be obtained from studies of neuronal oscillatory activity as an energy-efficient mechanism for temporal coordination of cognitive processes [26].

An increase of frontal midline *theta* rhythm (FMT) frequently accompanies such processes as nonspecific attention and WM [3,27–29]. The results of earlier studies often define FMT as the most plausible phenomenon reflecting an activation of central executive components of WM [30]. Several attempts to isolate central executive components from temporary storage components by including tasks requiring mental manipulations support hypothesis of the link between FMT and the executive control [31–33]. Several studies demonstrated the activation of fronto-parietal executive control system during retention in WM [34–36]. Moreover, some authors report increasing fronto-parietal synchronization with stronger engagement of central executive components [29]. Induced coupling of theta rhythm between frontal and parietal cortical regions by transcranial alternating current stimulation (tACS) resulted in improved visual WM performance, while the induced decoupling lead to WM deterioration [37].

Changes in *alpha* activity also show parametrical increase related to working memory load [38–40]. Increasing power of alpha rhythm is frequently interpreted as a mechanism for filtration and for suppression of the cortical areas irrelevant to the current task [40–42].

The role of *beta* activity in working memory processes is still not sufficiently investigated. Thus the activity particularly in the low beta band (~13-20 Hz) was found to increase during retention in WM [3,43,45]. A parametrical increase of low beta with the increasing of memory set size was also observed [3,43]. A comparison of retention condition with the conditions where participants were instructed to manipulate objects in WM showed that gradually increasing task complexity was related with a decrease of low beta activity [32].

Data of several studies suggest that the main contribution to individual differences in WM is made by the ability to control attention or executive control [46–48]. However, despite extensive research of WM in the recent 20 years, there is no clarity as regards the electrophysiological correlates mechanisms of individual differences in WM performance. The existing research (both general and differential psychological) have some limitations that restrict the possibility to explain the actual relationship between brain activity and WM performance.

First of all, most WM studies have used the n-back paradigm [1,2,49]. This kind of task engages multiple WM processes including retention of the stimuli set presented at the previous step, comparison between the first item of the memorized set and the new one, making decision about correctness of the comparison, and updating the content of WM. In this paradigm, it is difficult to clearly separate retention from the central executive components of WM.

Second, the level of difficulty of the task is usually moderate and thus does not present a big challenge for people with average WM abilities. There are studies dedicated to the investigation of EEG in WM tasks with several levels of difficulty [1,3,4,9]. In the studies mentioned above the number of steps did not exceed three (3-back) [1,2]. Some researchers applied other paradigms with gradually increasing difficulty of tasks for assessing WM performance [3,28,33]. But these paradigms either did not include any manipulation task [3,28], or their difficulty level was rather low [33].

Finally, the existing studies aimed to discover electrophysiological correlates of individual differences in WM were based on a sample size not exceeding 14 participants in each group [1,3,50]. An analysis of typical effect sizes indicates that at least twice larger groups would be necessary to reliably evaluate the differences between high- and low-performers.

In this paper we used highly demanding tasks which should give us the opportunity to distinguish EEG activity of individuals with different levels of WM performance. Additionally, using two types of tasks, which required either only retention of stimulus set or manipulation of content, we expected to reveal EEG correlates of temporary storage and central executive components of WM and to assess their contribution to individual differences.

The hypotheses of the study were as follows:

1. Motivated by the previous studies we expected significant relationships between WM performance and oscillatory activity in theta and alpha frequency bands;
2. Particularly, we supposed that frontal theta rhythm power is strongly related to the WM load;
3. We expected that storage components of working memory play less important role in individual differences than executive components. Specifically, we assumed that no individual differences would be found in the simple retention conditions;
4. Additionally, we hypothesized that the most challenging condition would best separate between low and high performers;

## Methods

### Participants

Due to a strong gender disproportion in the initial sample, only data of female participants were included into the present study. All participants were Russian native speakers. Furthermore, a subsequent analysis revealed five EEG records with an excessive amount of artefacts (i.e., less than 20 artifact-free epochs in at least one condition). Thus, 65 female participants (mean age = 20.92, SD=2.96) were included to the final sample. The participants had normal or corrected-to-normal vision and no history of neurological or mental diseases.

### Stimuli

Sets of Russian alphabet letters written in capital were used as stimuli. The letters had been selected randomly and had random order and no repetitions in the sets. Each trial consisted of 7 consecutive events. An analogue using Latin letters and English words is shown in Fig. 1.

**Fig. 1.**
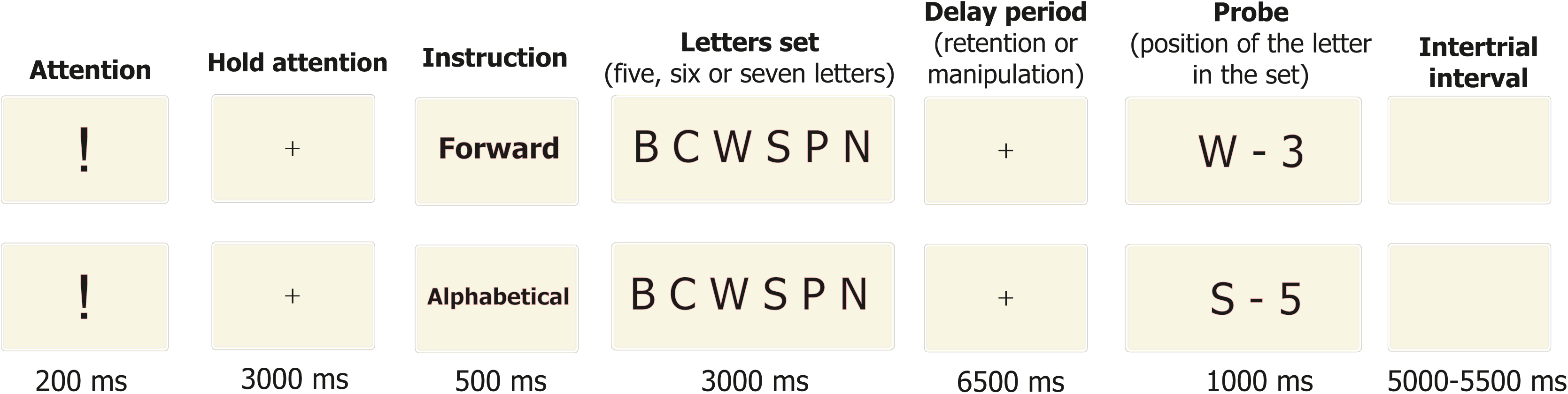
Examples of the trials.

A trial always began with an exclamation mark presented for 200 ms, which was followed by a fixation cross for 3000 ms. Participants were instructed to fixate the cross when it appeared in the center of the screen. At the next step the word “forward” or “alphabetical”, presented for 500 ms, instructed participants whether they would have to memorize the original set as it was presented (retention task) or to memorize it after mental recombination of the letters in the alphabetical order (manipulation task). After that, sets of 5, 6 or 7 letters were demonstrated for 3000 ms followed by a delay period where a fixation cross was demonstrated for 6500 ms. At the end of this delay period, a randomly chosen letter from the previously presented set appeared on the screen together with a digit that represented the serial number of this letter. The letter-digit combination was presented for 1000 ms. Participants were asked to press a specified button of a computer mouse if the presented letter had the corresponding serial number either in the original set (in the retention task), or in the set merging as a result of alphabetic recombination (in the manipulation task). The other mouse button had to be pressed if the serial number of the presented letter was incorrect. The two buttons were attributed to correct and wrong probes in a counterbalanced order. The probe was correct in 50 % of the trials, and the order of correct and incorrect probes was random. The next trial started after an interval that varied between 5000 and 5500 ms.

Thus, the experiment entailed six different conditions: memorizing 5, 6 or 7 letters in the alphabetical or forward order. Each condition had 20 consecutive trials. These six blocks with 20 trials were presented in a random order. A short practice block of 6 trials was given immediately before the main experiment.

During the experiment, the participants were seated in a comfortable armchair in front of a com puter screen in a dark room. Stimuli were presented in white color on a black background in the center of the screen by using PsyTask software (Mitsar Ltd.). The distance to the screen was 1 m and the size of the letters was 1.2 × 1.2°.

All participants were subdivided into two groups separated by the median of their mean performance across all tasks. The groups are referred to as high performance (HP; N = 32) and low performance (LP; N = 33) groups. The percentage of correct answers was used for behavioral data analysis. A repeated measures ANOVA with the between-subject factor Group (HP, LP) and the within-subject factors Task (retention, manipulation) and Load (5, 6, or 7 letters) was applied.

### EEG recording and analysis

The EEG was recorded from 19 electrodes arranged according to the 10-20 system using Mitsar-EEG-201 amplifier and referred to the average earlobe. Two additional electrodes were used for horizontal and vertical EOG. EEG data were acquired with 500 Hz sampling frequency, 0.16 Hz high pass filter and 70 Hz low pass filter.

Frequency bands for EEG analysis were defined using individual alpha frequency (IAF) as follows: theta = [IAF-6 Hz to IAF-2.5 Hz], alpha1 = [IAF-2.5 Hz to IAF], alpha2 = [IAF to IAF+2.5 Hz], beta1 = [IAF+2.5 Hz to 20 Hz], beta2 = [20 Hz to 30 Hz]. The IAF was determined on a 3 min EEG recorded at rest with eyes closed.

Segments of raw EEG recorded during the interval from 500 ms to 6500 ms of the delay period were analyzed. These segments were filtered between 0.5 and 30 Hz, and a 50-Hz notch filter was applied. The segments were subdivided into 2-second epochs. A fast Fourier transformation (FFT) was performed in each epoch. Ocular artefacts were corrected by using independent component analysis (ICA) followed by visual EEG inspection for remaining artefacts. These operations were performed in EEGlab toolbox. Spectral power densities for each frequency bands were calculated using Fieldtrip toolbox.

Spectral power data were statistically analyzed by using two independent mixed-design ANOVAs. The first analysis involved mean power values in four regions of interest (ROI): left (Fp1, F7, F3) and right (Fp2, F8, F4) anterior areas, left (T5, P3, O1) and right (T6, P4, O2) posterior areas. This analysis included a between-subject factor Group (HP, LP) and the within-subject factors Task (retention, manipulation), Load (5 versus 7 letters), Hemisphere (left, right) and Site (anterior, posterior). The second ANOVA of mean power values at the midline (Fz, Cz, Pz) analogous to the previous with factors Group, Task and Load was performed. All statistical calculations were performed by using SPSS package.

## Results

### Behavioral results

Participants performed with a general mean accuracy of 78.5±0.9%. Mean accuracies for each condition are shown in Fig. 2.

**Fig. 2.**
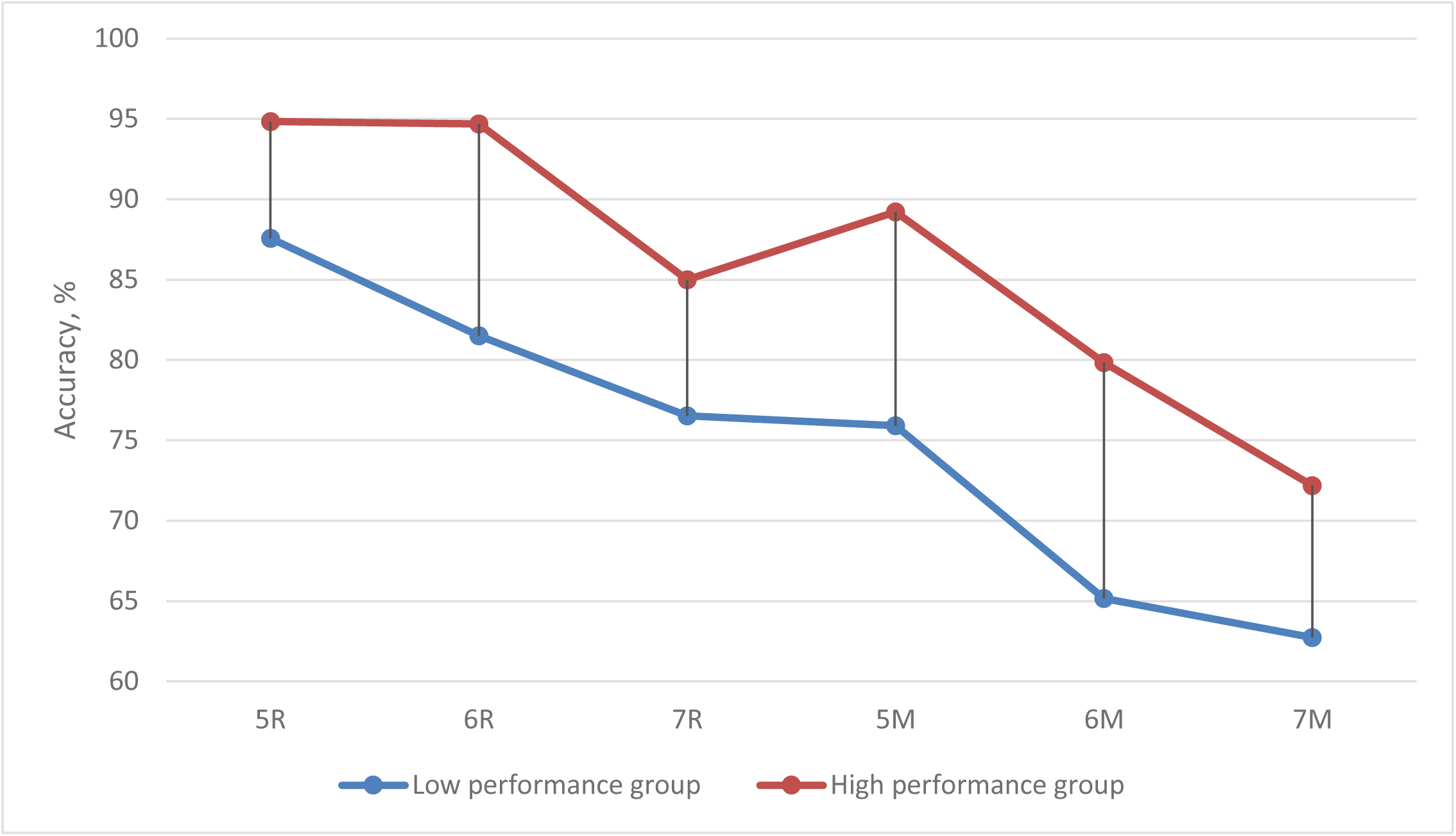
Mean accuracy in different WM tasks and conditions. Notes. 5R, 6R, 7R – memorizing 5, 6, or 7 letters in forward order (Retention condition); 5M, 6M, 7M – memorizing 5, 6, or 7 letters in alphabetical order (Manipulation condition).

The main effects of Task (F(1, 63) = 108.1, p < 0.0001, η2 = .632) and Load (F(2, 126) = 49.69, p < 0.0001, η2 = .441) as well as their interaction of the factors (F(2, 126) = 5.606, p = 0.005, η2 = .082) were obtained. A pairwise comparison between load levels separately for alphabetical and forward conditions showed highly significant differences (p<0.0001) for all pairs but two. First, there was no difference between the performance in 5- and 6-letter conditions in the forward order (p=0.191). Second, the differences were less pronounced in the comparison between 6 and 7 letters in the alphabetical order (p=0.011; not significant after Bonferroni correction). For this reason, and in order to avoid potential problem with sphericity in statistical measures, the 6-letters condition was excluded from the EEG analysis.

The mean performance accuracy in the high and low performance groups was 84.9±0,5% and 71.9±1,1%, respectively (F(1, 63) = 87.26, p < 0.0001, η2 = .581).

### Electrophysiological results

**Table 1.** Results of the ANOVA with the factors Task x Load x Hemisphere x Site x Group.

|  | *Theta* |  |  | *Alpha1* |  |  | *Alpha2* |  |  | *Beta1* |  |  | *Beta2* |  |  |
| --- | --- | --- | --- | --- | --- | --- | --- | --- | --- | --- | --- | --- | --- | --- | --- |
| | F | p | $\eta^2$ | F | p | $\eta^2$ | F | p | $\eta^2$ | F | p | $\eta^2$ | F | p | $\eta^2$ |
| *Task* | 3.631 | .061 | .054 | 9.694 | **.003** | .133 |  |  |  | 10.680 | **.002** | .145 |  |  |  |
| *Load* |  |  |  |  |  |  |  |  |  |  |  |  | 4.781 | **.033** | .071 |
| *Site* | 14.24 | **<.001** | .184 | 29.38 | **<.001** | .318 | 25.39 | **<.001** | .287 | 16.747 | **<.001** | .210 | 22.75 | **<.001** | .265 |
| *Hemisphere* | 7.712 | **.007** | .109 |  |  |  |  |  |  |  |  |  |  |  |  |
| *Group* |  |  |  |  |  |  | 6.143 | **.016** | .089 |  |  |  |  |  |  |
| *Site x Hemisphere* |  |  |  |  |  |  |  |  |  | 5.159 | **.027** | .076 |  |  |  |
| *Task x Group* |  |  |  | 4.763 | **.033** | .070 |  |  |  |  |  |  |  |  |  |
| *Task x Load* | 4.415 | **.040** | .065 |  |  |  |  |  |  | 6.376 | **.014** | .092 |  |  |  |
| *Task x Site* | 8.285 | **.005** | .116 | 5.620 | **.021** | .082 |  |  |  |  |  |  |  |  |  |
| *Task x Hemisphere* | 4.270 | **.043** | .063 |  |  |  |  |  |  |  |  |  |  |  |  |
| *Task x Site x Group* |  |  |  |  |  |  |  |  |  |  |  |  | 5.194 | **.026** | .076 |
| *Load x Site x Hemisphere* | 7.586 | **.008** | .107 | 5.363 | **.024** | .078 |  |  |  |  |  |  |  |  |  |
| *Task x Site x Hemisphere x Group* | 9.042 | **.004** | .126 |  |  |  |  |  |  | 5.131 | **.027** | .075 |  |  |  |

### General tendencies

#### Theta

The theta rhythm had lower power in anterior areas in comparison with posterior areas (main effect of Site, see Table 1 for this section). Also, the power was higher over the left than the right hemisphere (main effect of Hemisphere). Furthermore, the theta power decreased with the increasing WM load at all ROIs except the right anterior one (Load x Site x Hemisphere interaction).

Across the whole sample, the theta power tended to be higher in the manipulation task than in the retention task. As depicted in Fig. 3. this effect was more pronounced at anterior than posterior areas (Task x Site interaction) and also more pronounced over the left than the right hemisphere (Task x Hemisphere interaction).

The analysis of midline theta showed higher power in the manipulation task than in the retention task (main effect of Task, see Table 2). Increasing number of the presented letters from 5 to 7 yielded a decrease of theta power in the manipulation task but its increase in the retention task (Task x Load interaction). This interaction was, however, strongly modified by the between-subject factor, as described below in the Section Individual differences-Theta.

**Fig. 3.**
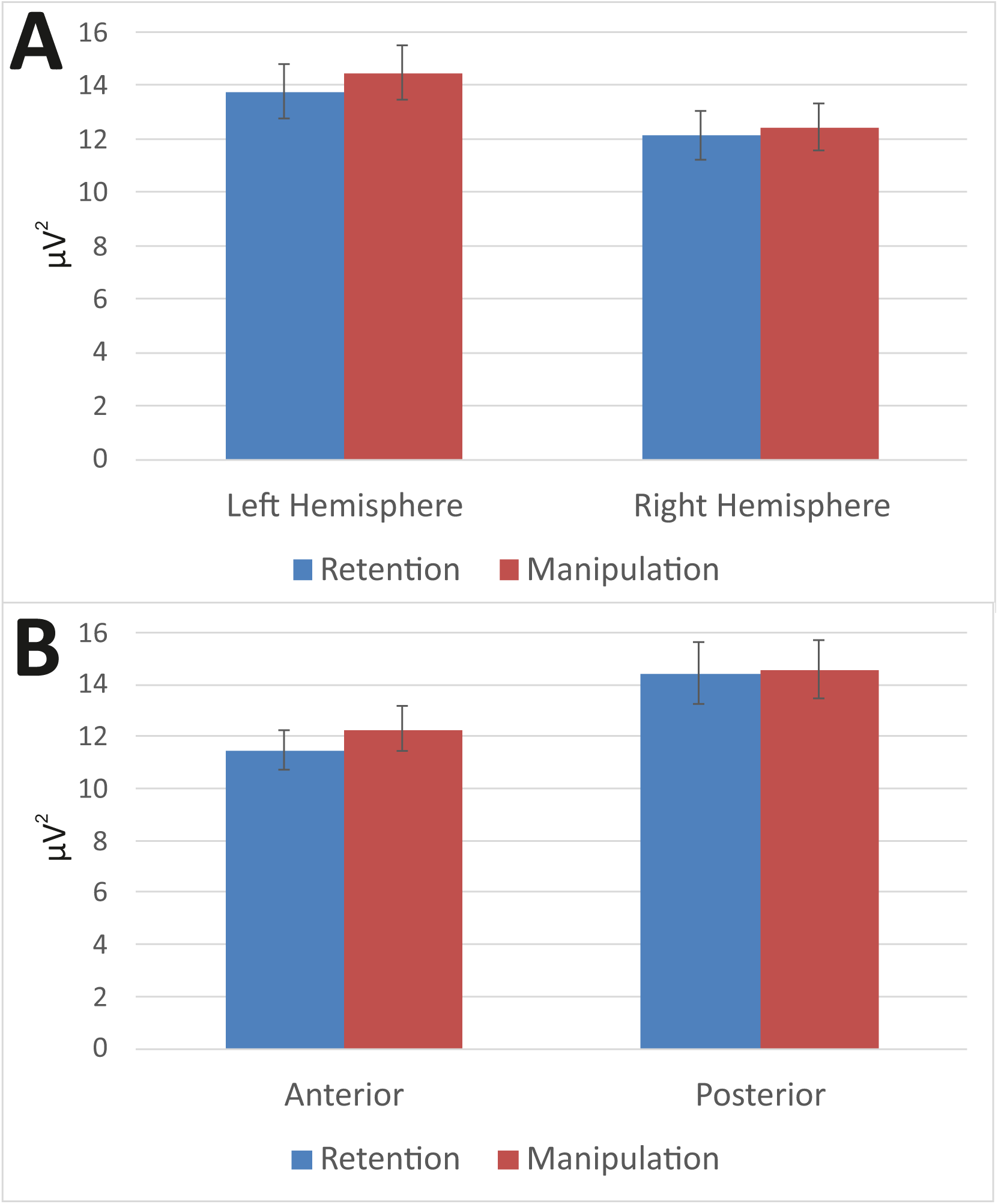
General tendencies of theta power for Retention and Manipulation tasks (A) over the Left and Right hemispheres and (B) in Anterior and Posterior areas. Error bars depict Standard Error of the Mean (SEM).

**Table 2.** Results of the ANOVA with the factors Task x Load x Group for midline sites.

| | | F | p | $\eta^2$ |
| --- | --- | --- | --- | --- |
| *Task* | *Theta* | 7.685 | **.007** | .109 |
|  | *Alpha1* | 6.526 | **.013** | .094 |
|  | *Beta1* | 20.427 | **<.001** | .245 |
|  | *Alpha1* | 4.715 | **.034** | .070 |
| *Load x Group* | *Theta* | 4.465 | **.039** | .066 |
| *Task x Group* | *Beta1* | 6.281 | **.015** | .091 |
| *Task x Load* | *Theta* | 5.462 | **.023** | .080 |
|  | *Beta1* | 5.895 | **.018** | .086 |

#### Alpha

As expected, alpha1 and alpha2 activity increased in the posterior direction (main effect of Site, see Table 1).

Alpha1 power was lower in the manipulation task than in the retention tasks (main effect of Task). This effect was larger at the posterior than anterior sites (Task x Site interaction).

Alpha1 activity was suppressed with increasing WM load in each ROI except the right posterior area where alpha1 power increased (Load x Site x Hemisphere interaction).

#### Beta1

Beta1 power was significantly lower in the anterior than posterior areas (main effect of Site), and lower on the left than right side (main effect of Hemisphere).

As can be seen in Fig. 4, beta1 power increased with the increasing WM load in the manipulation conditions but decreased in the retention conditions (Task x Load interaction). In general, the power was higher in the retention condition than in the manipulation condition (main effect of Task).

**Fig. 4.**
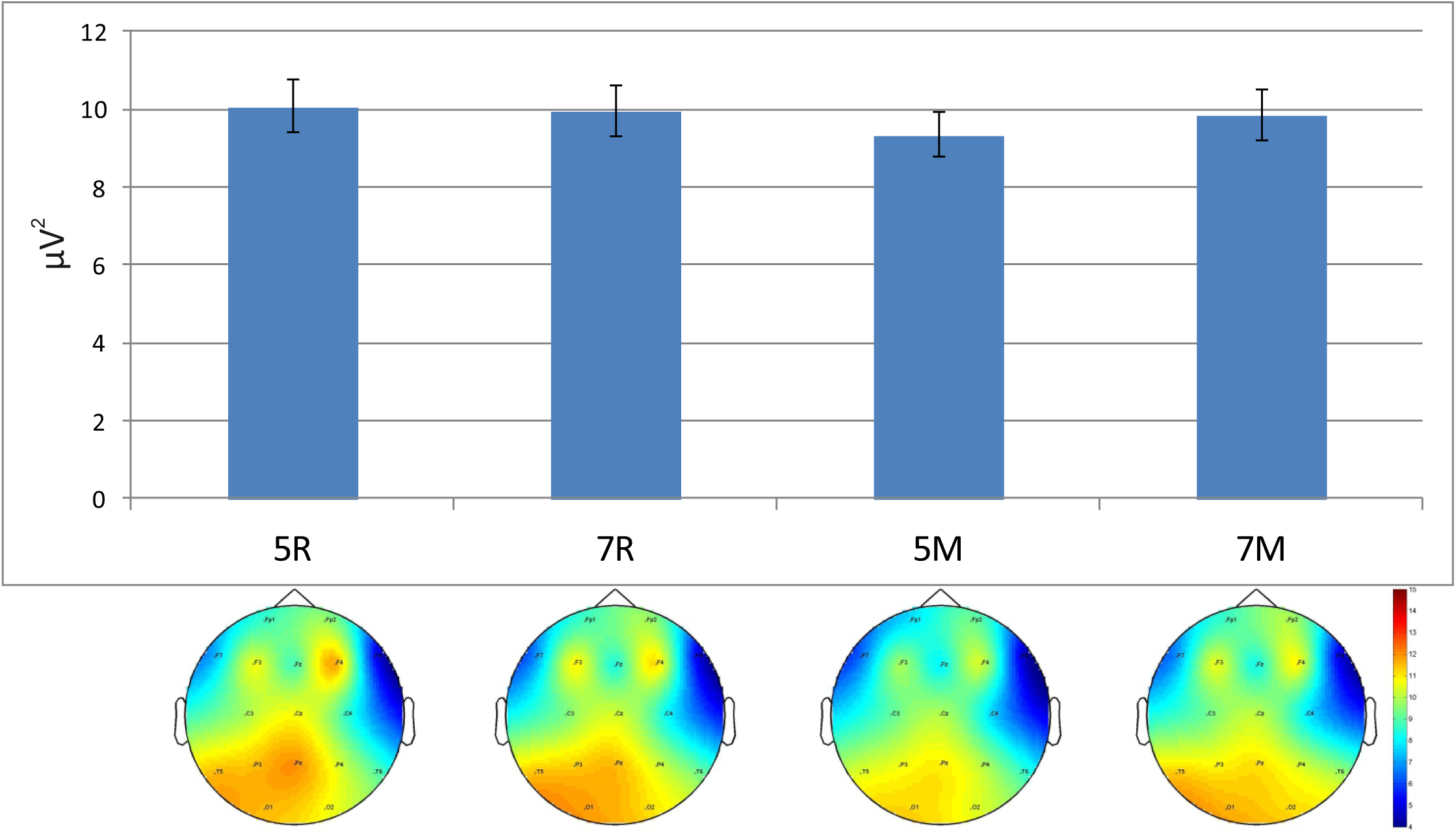
Beta1 power chart and corresponding topograms for Retention and Manipulation tasks. Error bars depict SEM.

#### Beta2

In contrast to beta1, beta2 power was significantly larger in the anterior than posterior areas (main effect of Site). Increasing WM load led to an increase in beta2 activity (main effect of Load).

### Individual differences

#### Theta

The analysis revealed a four-way interaction between Task, Site, Hemisphere and Group. Additional separate analyses in groups were performed. In the HP group we observed a larger theta power in the manipulation condition than in the retention condition, and the magnitude of this effect was the highest in the left anterior area (Task x Site x Hemisphere interaction (F(1, 31) =7.605, p = 0.01, *η^2^* = .197). No significant effects were found in the LP group.

An ANOVA performed on midline electrodes revealed opposite load dependent changes of the midline theta power in the HP and LP groups. As depicted in Fig. 5, an increase of the number of letters from 5 to 7 was associated with an increase of theta activity in the former group but its decrease in the latter (Load x Group interaction, see Table 1). Fig. 5 shows that the significant Load x Task interaction for the entire sample, described above in Section General tendencies-Theta, is actually produced by the dramatic decrement of the theta power in the most demanding condition (manipulation task, high WM load) in the LP group. Similarly, the triple interaction Load x Site x Hemisphere for the entire sample does not really characterize the entire sample but, like the Load x Task interaction, can be attributed to a disproportionately strong influence of the LP group.

**Fig. 5.**
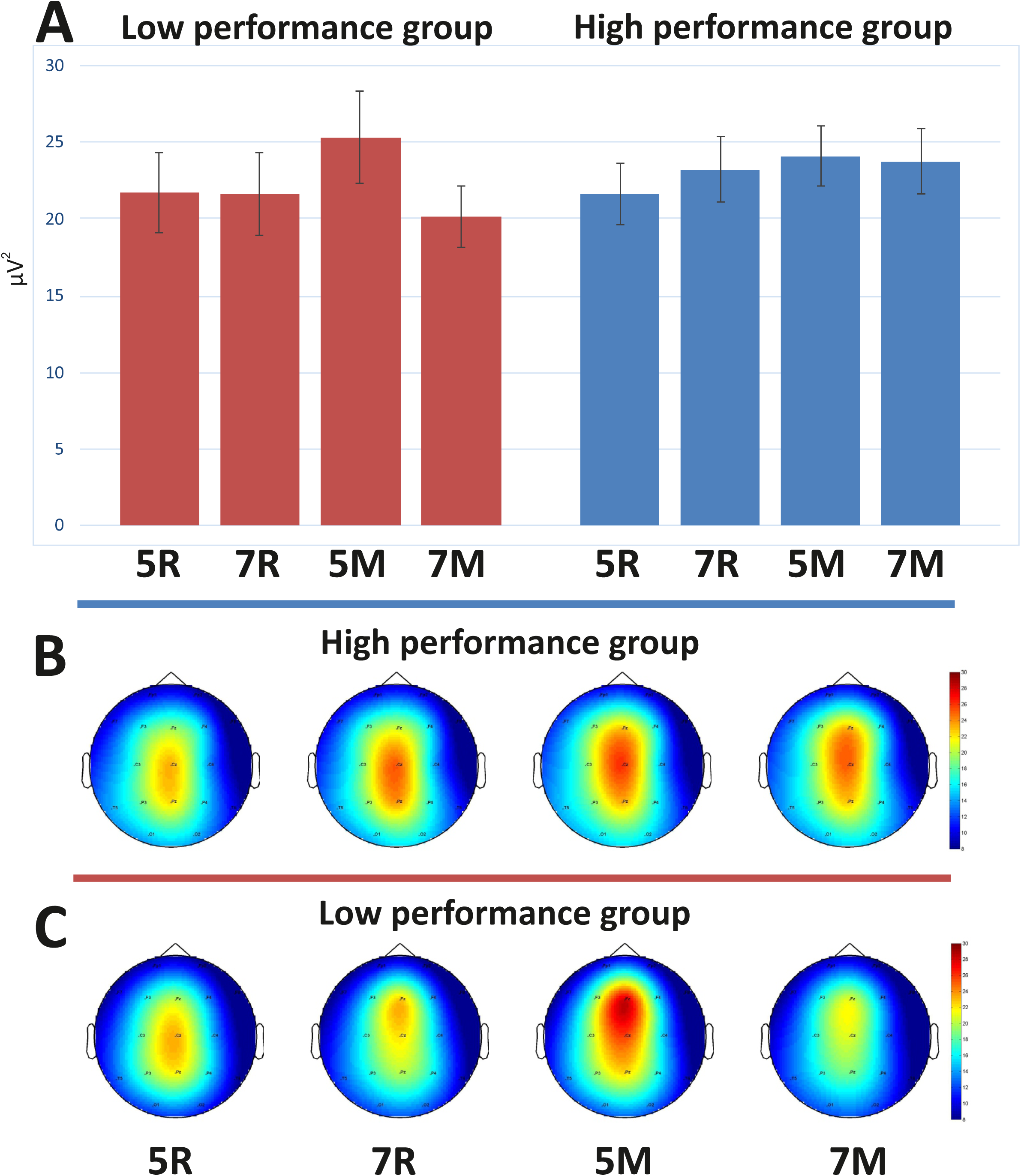
(A) Midline theta power for four WM tasks and (B, C) corresponding topograms in two groups. Notes: 5R, 7R – 5 and 7 letters Retention conditions; 5M, 7M –5 and 7 letters Manipulation conditions. Error bars depict SEM.

#### Alpha

As can be seen in Fig. 6, the suppression of the alpha1 power in the manipulation task relative to the retention task was stronger in the HP than the LP group (Task x Group interaction). Alpha2 was generally stronger in the HP than the LP group (main effect of Group).

**Fig. 6.**
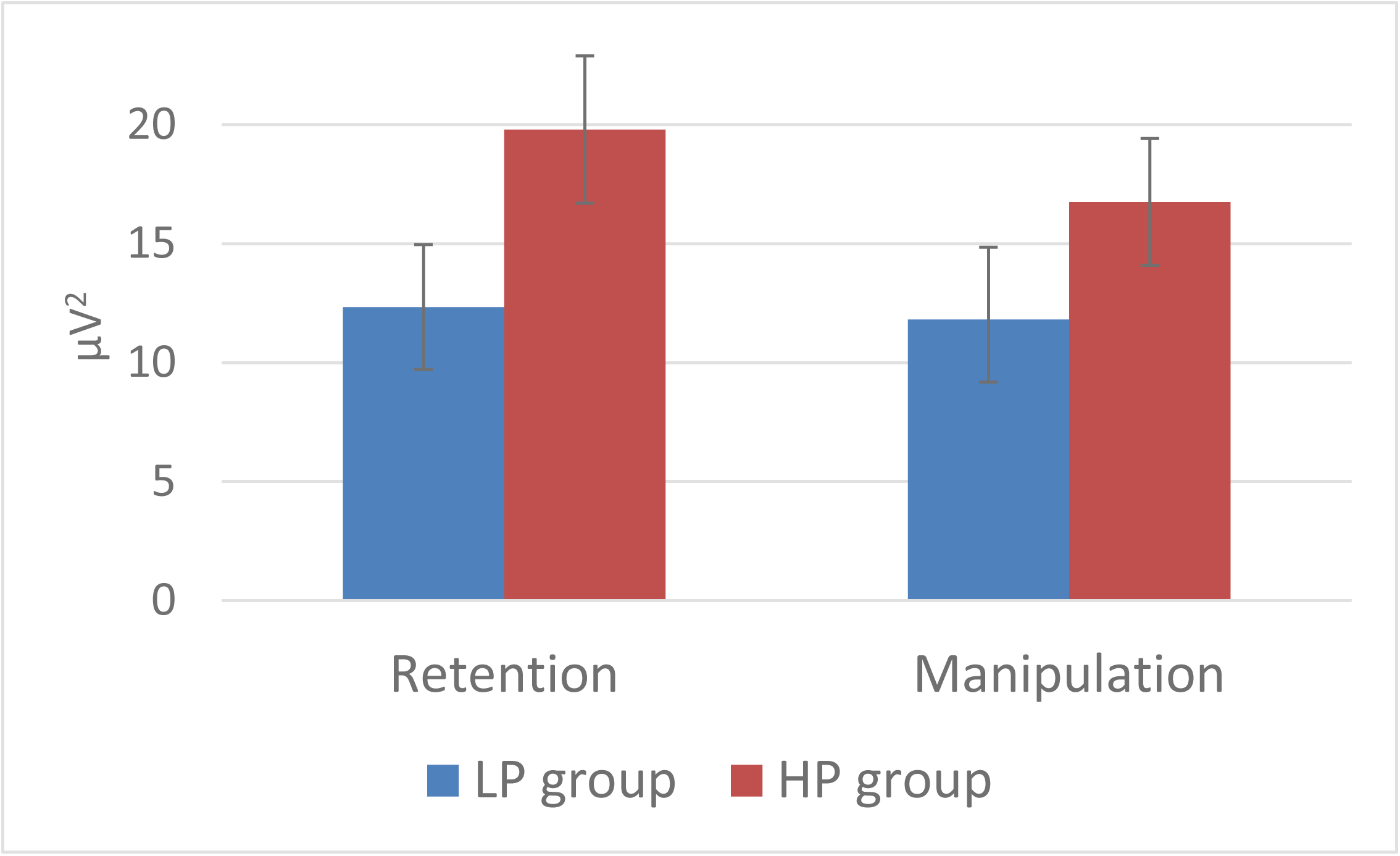
Alpha1 power for Retention and Manipulation tasks in Low and High performance groups. Error bars depict SEM.

#### Beta1

A significant four-way interaction Task x Site x Hemisphere x Group was obtained and further analyzed for groups and for electrode sites. The first ANOVA yielded a significant Task x Site x Hemisphere interaction (F(1, 31) =6.471, p < 0 .05, *η^2^* = .131) only in the HP group, indicating that the decrease of the beta1 power from the retention task to the manipulation task was more pronounced in the left posterior and the right anterior ROIs. No such effects were observed in the LP group.

The second ANOVA revealed a significant Task x Group interaction in the left posterior ROI (F(1, 31) = 5,953, p < 0.05, *η^2^* = .086), indicating task dependent changes of beta1 power at the left posterior area in the HP group (see Fig. 7).

**Fig. 7.**
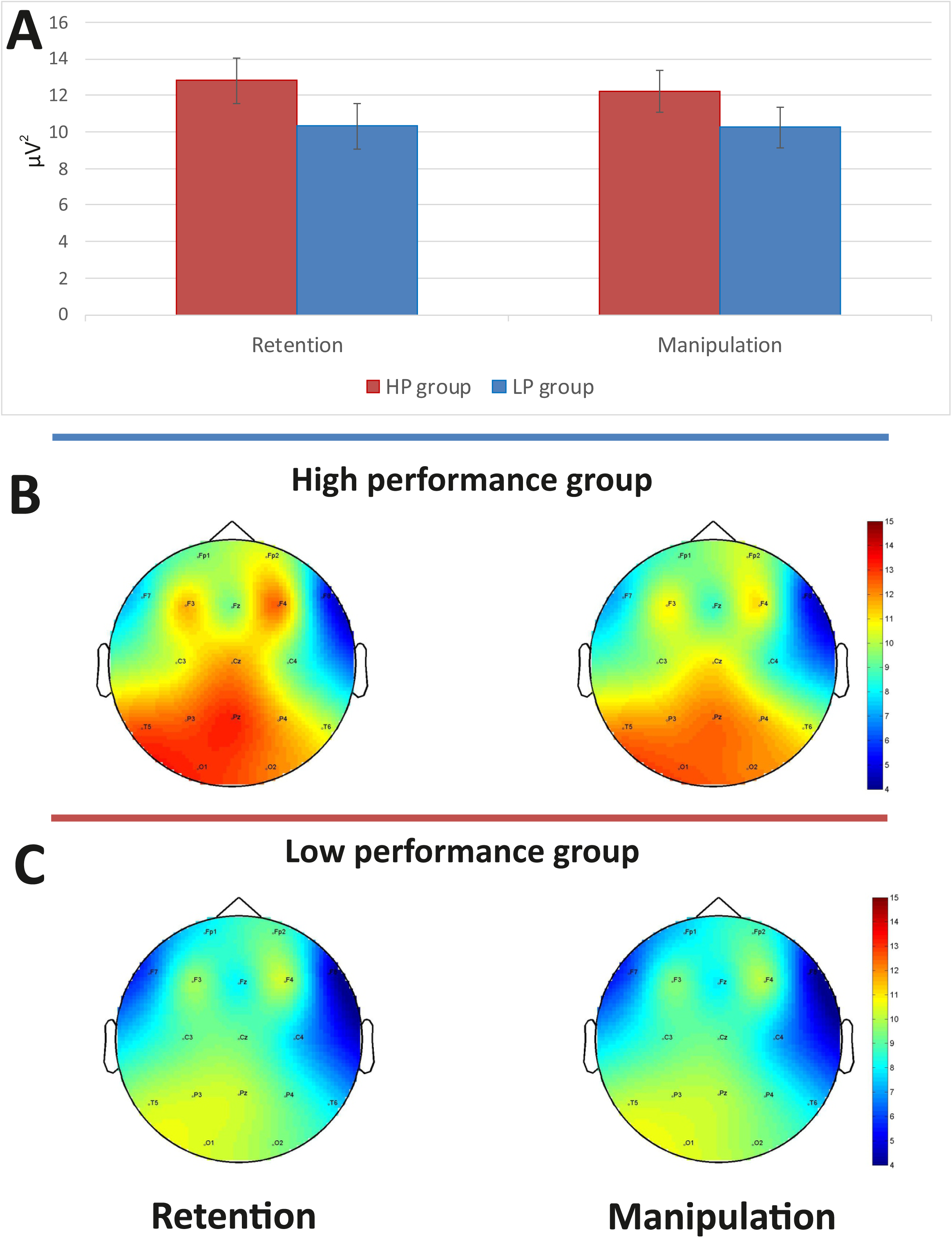
(A) Beta1 power in the left posterior area for Retention and Manipulation tasks and corresponding topograms (B) in Low performance (LP) and (C) High performance (HP) groups. Error bars depict SEM Fig. 8 Beta2 power for Retention and Manipulation tasks in Low and High performance groups in Anterior and Posterior areas. Error bars depict SEM.

#### Beta2

The significant Task x Site x Group interaction (see Table 1) indicates opposite task- and location-related changes in the two groups. The LP group showed higher beta2 activity in the manipulation task at anterior areas but in the retention task at posterior areas, while the opposite held true for the HP group (Fig. 8).

**Fig. 8.**
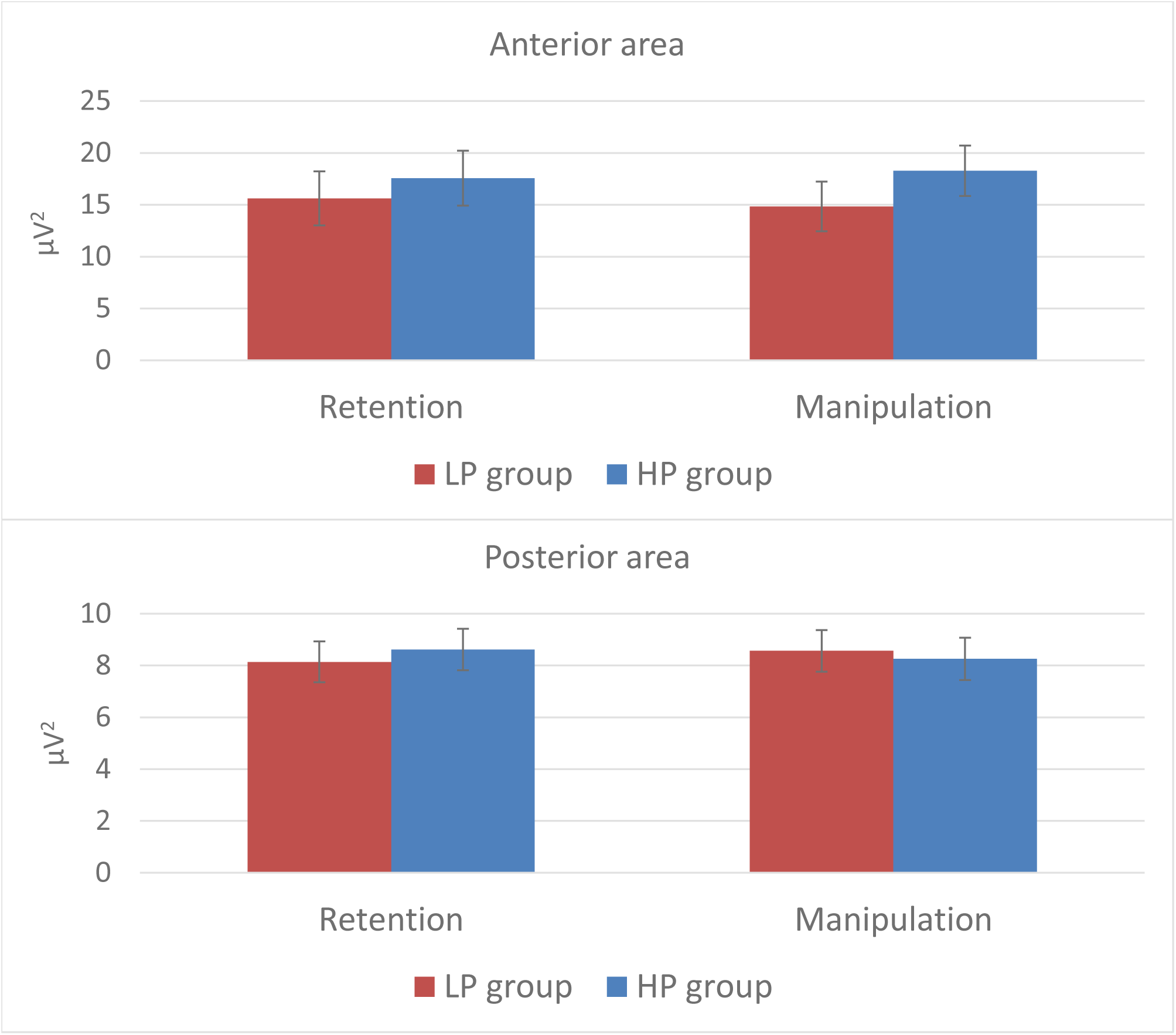
Beta2 power for Retention and Manipulation tasks in Low and High performance groups in Anterior and Posterior areas. Error bars depict SEM.

## Discussion

### General tendencies

#### Theta and central executive components of WM

The current study found that increasing WM task complexity and executive control demand were associated with the increase of the frontal theta activity. Increasing theta power in midline and frontal areas during mental manipulations in contrast to the mere retention of memory content is in line with numerous data indicating positive relationships between FMT and cognitive load [3,9–11,28]. Moreover, an increase of FMT in manipulaton tasks as compared with retention tasks was also found in studies whose design was similar to the present one [31–33,52].

In addition, the link between FMT and the activation of the anterior cingulate cortex (ACC) and the medial prefrontal cortex (mPFC) was repeatedly proven by simultaneous EEG-fMRI recordings as well as by direct electrophysiological recordings in monkeys [53–56]. The ACC and the mPFC are active during memory processes, WM performance, and executive control [57–59].

We assume that the increment of FMT (supposedly indicating the activation of the ACC) with increasing WM demands is related to increasing involvement of executive processes. However, it should be noted that FMT reflects not pure memory processes per se but more likely the allocation of cortical resources depending on features of the task [3,59,60]. One may speculate that increasing demands for executive control during manipulation of information in WM engage a widely distributed network whose main components are the prefrontal cortex and the ACC.

#### Alpha1 and the storage components of WM

As compared with the retention condition, manipulation of stimuli in WM was associated with distributed suppression of alpha1 activity. There is an empirically well supported hypothesis that desynchronization of low alpha is a nonspecific cortical response that can be observed during various cognitive operations [9,38,61] including maintaining information in WM [53,62,63]. In addition to this non-specificity model, however, more specific hypotheses about the dynamics of alpha exist. Thus, the alpha synchronization in posterior areas during the maintenance of actual information may reflect active inhibition to protect these areas from reorienting to new irrelevant information processing [28,42]. It is plausible that the temporary storage components of WM play the key role in a successful maintenance of 7 letters relative to 5 letters. It might be suggested that when the volume of information maintained in the temporary storage approaches the putative capacity limit (7+-2) the central executive should actively inhibit irrelevant information. The observed asymmetry of alpha1 power at the posterior area agrees with the previous studies of WM and short-term memory [9,28,54,64,65].

#### Beta1 and manipulation of information in WM

Task-related decrement of beta1 power found in this study was quite similar to the effect reported by Berger et al. [32] who also compared manipulation versus retention conditions. This effect may be explained by the conception of Engel & Fries [66] that, applied to the present experiment, suggests that the decrease of beta1 power takes place during updating or manipulating information in WM as well as during retrieval of information from long term memory and encoding it in WM. The desynchronization of the beta1 rhythm can be attributed to the sequential updating of the WM content during mental alphabetizing of the letters. This process also involves addressing the long term memory where the alphabet is stored.

Load-dependent changes in beta1 power were observed only in the manipulation condition. We hypothesize that manipulation is underpinned by two independent temporal buffers: the first one is the final storage for modified items after the manipulations, whereas the second one serves as a workspace for the remaining to-be-modified items. Perhaps, there are even two different beta1 rhythms that overlap in frequency but reflect different sub-processes in WM [67]. The first rhythm supports the activity of the first buffer (“store”), and the second rhythm, that of the second buffer (“workspace”). Synchronization of the former maintains the active state of the engram and protects it from irrelevant information. Weiss & Rappelsberger [68] demonstrated a gradual increase of beta1 activity in response to sequential filling of WM by words. Research conducted by Leiberg, Lutzenberger, & Kaiser [69] also showed a load-dependent increase of beta1 activity. At the same time desynchronization of the other beta1 rhythm reflects the retrieval from long term memory and encoding to WM. In other words, desynchronization of the latter beta1 rhythm reflects manipulations of objects in “workspace” for their subsequent transfer to “store”.

Our hypothesis also entails that the lack of beta1 desynchronization during the encoding process indicates a disruption of memory formation. Recently, Hanslmayr, Matuschek, & Fellner [70] found a negative effect of transcranial magnetic stimulation (TMS) of the left inferior frontal gyrus at beta1 frequency (18.7 Hz) on memory performance in a word-list learning task. Furthermore, a study [44] performed on monkeys demonstrated desynchronization of beta activity during updating of WM content but synchronization of beta activity during retention.

Probably, in the retention condition the “workspace” buffer is minimally involved. It may work at the beginning of the delay period when sequentially and quickly presented information is encoded. Thus Zanto & Gazzaley [71] found the desynchronization of beta1 rhythm during the first 1250 ms of the 4-s delay period but the synchronization from 1500 ms to the end of the delay. In the current study, the delay periods during maintaining and manipulation of 5 and 7 letters could be different due to a longer presentation time (3 seconds). Therefore, the recombination of 5 letters to the alphabetical order could already start during stimulus presentation and continue only in the “workspace” buffer without addressing the “store” buffer. When the recombination process is finished, the result is transferred to the “store” buffer and kept there until the probe is presented. The “store” buffer in this case prevents possible interference of other stimuli and maintains the actual state of the engram until the moment when its content is requested. When a longer stimulus set is memorized (i.e., 7 letters) a plausible strategy is to memorize the initial letters set and to transfer it into the “store” buffer. If this strategy is used, recombination may start after the stimuli have disappeared from the screen. During this period, both buffers are actively involved: the “store” buffer is keeping the initial set, while recombination is carried out in the “workspace” buffer. When the recombination is finished the information transfers to the “store” and updates its content. This assumed information return to, and updating of, the “store” buffer would explain the increase of beta1 power from 5- to 7-letter condition in the manipulation task.

#### Beta2 and amount of information in WM

Beta2 power increased with the increasing WM load and did not significantly depend on the type of the task.

Dissociations between the lower (13-20 Hz) and upper (25-30 Hz) beta were demonstrated earlier in a study of Shahin, Picton, & Miller [72]. The authors concluded that the increment of the upper beta may reflect maintaining verbal stimuli in auditory memory. The maintenance of stimuli in WM was also suggested to cause synchronization of beta2 (~20-30 Hz) in two different tasks [73,74]. Spitzer et al. [74] assumed that the upper beta activity is directly related to the quantity of supramodal abstract information. The significant effect of Load on beta2 power found in the present study is in line with this interpretation.

### Individual differences

#### Theta

The task-related increment of theta power in the left anterior area was found only in the HP group. This may be related to more effective manipulations supported by the hippocampus and language cortex. Previous WM studies demonstrated a relationship between the hippocampal activity and the theta rhythm [75,76]. In animal studies, the synchronized activity of the prefrontal cortex and the hippocampus crucially determined the accuracy in WM tasks [77,78]. The prefrontal cortex is hypothesized to be supported under excessive WM load by the medial temporal lobe related to long term memory [79–81].

The activation of the left prefrontal cortex including the inferior frontal gyrus (IFG) and Broca's area was found in verbal tasks during executive processes functions [55,82,83]. Simultaneous EEG / fMRI recording in a modified Sternberg task revealed a load-dependent increase of left IFG activation and the theta rhythm [55]. Similar results were obtained by Chee & Choo [84] in a WM task. We suppose that the left-hemispheric accentuation of the theta rhythm represents more effective information exchange between short- and long-term memory storage in the HP group.

Group differences were not only task-dependent but also load-dependent. The HP group demonstrated a gradual increase of theta power at midline, reaching its peak in the most demanding condition: manipulation task with 7 letters. In contrast, the LP group exhibited a sharp drop of theta power in this condition after a maximum in the condition of moderate difficulty: manipulation with 5 letters. Since previous studies of EEG correlates of individual differences in WM were limited to moderate difficulty, we can state that our findings are fully consistent with the previous ones, where the theta activity always increased with memory load [1,3,4,9–11]. However, the most difficult task resulted in a more complex change of theta activity that has not been observed so far.

One may speculate that reaching the individual’s WM capacity limit is accompanied by a crucial deficit of attentional resources. Post-experimental reports suggest that most participants formulated their task as “to remember all letters if possible”, but possibly, some LP participants in the most difficult condition changed the task to “to remember at least some letters”. Alternatively, some subjects may have switched strategy to “remember the first few letters with regard to position” in the forward task and the “first few letters with regard to alphabetical order” in the alphabetical task. This post-hoc hypothesis was tested by an analysis of behavioral results with regards to the position of the probe letter. The factor Position was taken with 2 levels (the first two versus the last 2 letters for 5-letters conditions, or the first three versus the last 3 letters for 7-letters conditions). We found two significant interactions between Position and Group: Position x Group (F(1, 63) = 6.022, p = 0.017, *η^2^* = .087) and Position x Task x Load x Group (F(1, 63) = 3.183, p = 0.045, *η^2^* = .048). Unfortunately, due to the post-hoc nature of this effect we could not perform the EEG analysis with the factor Position, because we did not have a sufficient statistical power for this unplanned comparison.

Another explanation might be the loss of motivation in LP participants in the most challenging condition. This hypothesis, however, would predict a particularly poor performance of LP participants in the manipulation task with 7 letters. This disagrees with the observed data indicating nearly equal performance differences between LP and HP participants in all conditions (see Fig. 5). From our point of view, the strategy change hypothesis can better integrate this fact that the loss-of-motivation hypothesis.

Also Jaeggi et al. [2] came on the basis of their fMRI study to the same conclusion concerning the suboptimal strategies used by LP subjects in WM tasks. In that study, LP participants showed a positive correlation between task complexity and the amount of the broad activation in the frontal cortex. Obviously, the most challenging condition leads to the widely distributed engagement of the prefrontal cortex and results in the lack of neural resources for activation of the ACC necessary for the executive control of WM.

#### Alpha

In the development of the cortical idling hypothesis, [85] proposed that the increasing alpha activity during cognitive processing is related to the allocation of attentional resources by inhibition of the cortical areas irrelevant to the current task [42,86,87]. In this context alpha rhythm plays the role of an information flow filter.

It is well known that WM is one of the main components of general intelligence [88,89]. Accordingly, the degree of alpha desynchronization in semantic memory task is positively related to intelligence [90]. Similar correlations between IQ and alpha power were observed in the resting state [91,92]. We suppose that stronger alpha power may reflect a higher level of readiness to perceive relevant information. Therefore, HP individuals have potentially more resourceful visual cortex and manage the tasks better [61].

#### Beta1

The main result was a stronger desynchronization of beta1 rhythm in the HP group in the manipulation condition in the left posterior area. An important role of the superior parietal cortex in flexible redistribution of attentional resources was demonstrated in several studies [93–96]. In terms of the two-buffer model (see above, Section General tendencies-Beta1), one may suggest that HP individuals are better able to shift their attention between the store of the originally presented set and the workspace where they work with the symbols. This might allow them to perform manipulations in the “workspace” buffer not spending too much resources for maintaining information in the “store” buffer.

#### Beta2

In the manipulation task, beta2 power increased in the HP group in the anterior areas, but in the LP group in the posterior areas. As we do not know any comparable data in the literature, only a very preliminary explanation can be proposed. Beta2 is the EEG index that may most simply be designated as “activation”. We believe, therefore, that changes in beta2 activity are not related to mental processes as such, but rather to the general volume of information necessarily used in these processes. This volume is expected to be larger in the manipulation task than in the retention task because during manipulation one has to work with at least two stimulus sets: the one that should be manipulated with and the one that results from the manipulation. The increase of frontal activity in HP participants may, therefore, reflect their ability to process a larger amount of information, whereas the heightened activity of sensory regions in LP subjects appears to reflect their need to frequently address the original stimulus set.

#### General discussion

In general, the obtained results allow us to make several claims about possible factors contributing, at the individual level, to effective verbal WM performance:

firstly, a higher state of readiness to process relevant and to inhibit irrelevant information and related larger alpha power;
secondly, stronger engagement of the left prefrontal cortex and the hippocampus; this factor can underlie efficient maintaining and manipulating information in WM due to a fast exchange of information between long term and working memory;
thirdly, an energy efficient strategy for distribution of frontal resources in order to maintain the necessary level of activity of the ACC;
finally, activation of the ACC and the related executive functions is decisive for successful manipulations of content in WM, simultaneous maintaining information about initial properties of stimuli and efficiently shifting attention between these cognitive operations.

## Limitations

We have to acknowledge at least two limitations of the present study. Firstly, the results may be affected by the homogeneity of the sample in respect to gender (i.e., females). A gender based analysis will be the matter of a subsequent report Secondly, our putative explanation hypotheses suggested in the Discussion above have neuroanatomical implications, i.e., they presume the activity of certain brain structures such as the ACC. To test these hypotheses, a larger number of electrodes should be used in future studies, which will allow a more precise assessment of the spatial distribution of the obtained effects.

## Conclusions

1. In accordance with many previous studies, we expected to find significant WM-related changes in alpha and theta frequency bands. This hypothesis was only partially supported by the data. Significant effects were found in all analyzed frequency bands from theta to high beta, indicating that our knowledge about the neural basis of WM is not comprehensive.
2. The hypothesis about a strong participation of the frontal theta rhythm in WM processes was confirmed. The novel finding was, however, different dynamics of frontal theta in HP and LP groups.
3. When starting the study, we believed that some important findings can have been missed in the previous experiments because they used only tasks of low to average difficulty. Therefore, we predicted important intergroup variation in EEG pattern in the most challenging condition. This prediction was confirmed. The most pronounced differences between individuals with high and low WM performance, in terms of the oscillatory activity in several frequency ranges, were observed in the manipulation task with 7 letters, which is a very difficult condition that for many individuals might exceed their limits. Particularly, this condition resulted in a more complex change of theta activity than just an increase with WM load, which has not been observed so far. Including greater variety of experimental conditions and groups to the WM research agenda seems beneficial.
4. Finally, we expected a stronger effect of executive WM components as compared with storage components. The data put this hypothesis in question. Firstly, the difference in performance between LP and HP participants was nearly equal in retention (weak executive control demands) and manipulation (much higher executive control demands) conditions. Secondly, task and site dependent group differences were found in each explored frequency bands including anterior theta and posterior alpha activity. In some studies these two responses were interpreted as reflections of executive and storage components of WM, respectively [5,6]. Although there is an alternative interpretation on the basis of cross-frequency coupling [7,8], all these observations together may indicate that the two components of WM are equally important for WM performance at the individual level. More studies are needed to clarify this issue

## Declarations

## Abbreviations

WM: working memory
HP: high performance
LP: low performance
tACS: transcranial alternating current stimulation
FMT: frontal midline theta rhythm
EEG: electroencephalography
EOG: electrooculography
IAF: individual alpha frequency
FFT: fast Fourier transformation
ICA: independent component analysis
ROI: region of interest
SEM: standard error of the mean
IFG: inferior frontal gyrus
fMRI: functional magnetic resonance imaging
ACC: anterior cingulate cortex
IQ: intelligence quotient
ANOVA: analysis of variance

## Ethics approval and consent to participate

Informed consent was obtained from all subjects prior to the study. The study was approved by the Ural Federal University Ethics Committee.

## Consent for publication

Not applicable.

## Availability of data and materials

The datasets analyzed during the current study is available from the corresponding author on reasonable request.

## Authors’ contributions

**YGP**: conceived of the study, designed the experimental paradigm, carried out the recording of the data, performed the statistical analysis and drafted the manuscript

**BK**: contributed to the discussion, and to the preparation of the manuscript

All authors read and approved the final manuscript.

## Acknowledgements

Not applicable.

